# Cross-decoding reveals shared brain activity patterns between saccadic eye-movements and semantic processing of implicitly spatial words

**DOI:** 10.1101/415596

**Authors:** Markus Ostarek, Jeroen van Paridon, Falk Huettig

## Abstract

Processing words with referents that are typically observed up or down in space (up/down words) influences the subsequent identification of visual targets in congruent locations. Eye-tracking studies have shown that up/down word comprehension shortens launch times of subsequent saccades to congruent locations and modulates concurrent saccade trajectories. This can be explained by a task-dependent interaction of semantic processing and oculomotor programs or by a direct recruitment of direction-specific processes in oculomotor and spatial systems as part of semantic processing. To test the latter possibility, we conducted a functional magnetic resonance imaging experiment and used multi-voxel pattern analysis to assess 1) whether the typical location of word referents can be decoded from the fronto-parietal spatial network and 2) whether activity patterns are shared between up/down words and up/down saccadic eye movements. In line with these hypotheses, significant decoding of up vs. down words and cross-decoding between up/down saccades and up/down words were observed in the frontal eye field region in the superior frontal sulcus and the inferior parietal lobule. Beyond these spatial attention areas, typical location of word referents could be decoded from a set of occipital, temporal, and frontal areas, indicating that interactions between high-level regions typically implicated with lexical-semantic processing and spatial/oculomotor regions constitute the neural basis for access to spatial aspects of word meanings.

## Introduction

The capacity of the adult human brain to virtually effortlessly derive abstract meaning from language input poses an enormous challenge for contemporary cognitive neuroscience. One influential proposal has been that language comprehension is made possible by grounding it in the sensory and motor systems of the brain (Barsalou, 1999; Barsalou, Simmons, Barbey, & Wilson, 2003). According to this proposal, conceptual processing of spoken and written words involves a recycling process: It necessitates the re-recruitment of sensory-motor processes that were consistently activated during previous encounters with the ‘real world’ referents the words denote. In the past decades, our understanding of the cognitive and neural mechanisms underlying conceptual processing has advanced considerably (Ralph, Jefferies, Patterson, & Rogers, 2017). Recent evidence suggests that the brain’s solution to the complex problem of knowledge storage and conceptual processing has indeed not been to evolve a new anatomically defined self-sufficient semantic module but to employ a widely distributed network that seems to encompass most of the cerebral cortex (Huth, de Heer, Griffiths, Theunissen, & Gallant, 2016; Mitchell et al., 2008).

When it comes to the question of how conceptual knowledge is represented in neural codes, most studies to-date have targeted concepts with concrete referents that are straightforwardly described in terms of their sensory-motor features. For instance, action words have often been found to activate motor areas implicated with executing or planning movements required for the action (Hauk, Johnsrude, & Pulvermüller, 2004; Vukovic, Feurra, Shpektor, Myachykov, & Shtyrov, 2017). Similarly, words referring to objects with salient visual characteristics recruit the corresponding visual processes/areas (shape: Correia et al., 2014; Lewis & Poeppel, 2014; Ostarek & Huettig, 2017b, 2017a; color: Simmons et al., 2007; speed of motion: van Dam, Speed, Lai, Vigliocco, & Desai, 2017). Recent studies using multivariate pattern analyses (MVPA) suggest that activation patterns in areas associated with high level semantic processing (Binder, Desai, Graves, & Conant, 2009) at least partly reflect conjunctive coding of multiple sensory-motor attributes (Fernandino, Humphries, Conant, Seidenberg, & Binder, 2016; Fernandino et al., 2015), allowing encoding models based solely on sensory-motor attributes to predict voxel-wise activity patterns in response to concrete - but not abstract - words (Fernandino et al., 2016). Thus, the emerging picture is a hierarchical organization of semantic knowledge derived from experiential sensory-motor experience (Fernandino et al., 2016) that may culminate in holistic multimodal concept-level neural codes in high-level areas (Binder & Desai, 2011; Coutanche & Thompson-Schill, 2014; L. Fernandino et al., 2016; Ralph et al., 2017).

We thus have a growing understanding of both the architecture and the processes related to item-dependent concepts with referents that map onto a set of sensory-motor features. However, it is still largely unclear what neurocognitive mechanisms are recruited for item-*independent* conceptual dimensions. These include abstract concepts, such as *justice*, which do not have tangible referents with consistent physical properties, and spatial dimensions, such as *up* and *down*. Researchers have so far struggled to explain how this important aspect of our conceptual knowledge is enabled on the cognitive and neural level. At least partly, this is due to the fact that researchers have thus far failed to get a good grasp on the kinds of relevant features because of the lacking consistency of features across instantiations of most abstract concepts (Borghi et al., 2017).

Here, we tested the hypothesis that the dorsal pathway provides semantic information about vertical space, a dimension largely orthogonal to features of individual items (Ralph et al., 2017). More specifically, we examined the processing of words that differ in the vertical spatial location where their referents are typically observed (e.g., *bird* vs. *foot*, henceforth ‘up/down words’) in order to address how the abstract features ‘up’ and ‘down’ are reflected in neural population codes using multi-voxel pattern analysis (MVPA). The vertical dimension is not only abstract because it is item-independent, but also because it can only be indirectly experienced by virtue of being defined through varying reference frames. This makes the dimension a great testbed for the scope of the conceptual grounding hypothesis; if a dimension that is in principle an ideal candidate for abstract representation in a high-level system is nevertheless anchored in sensory mechanisms, this would indicate that grounding may be the primary route to meaning.

If this were the case, it would make sense to capitalize on to the most reliable way of engaging with up/down items. The best candidate is the cortical substrate for vertical looking behaviour, primarily the one related to saccadic eye movements. This is because looking behaviour is a prime attentional gate keeper that largely determines which entities we consciously perceive. Therefore, in a typical situation in which you perceive a bird (perhaps because somebody says “Look at that big bird”) you move your eyes up to look at it.

Numerous studies have shown that processing up/down words can influence performance at visual tasks involving compatible vs. incompatible location (Estes, Verges, & Adelman, 2015; Estes, Verges, & Barsalou, 2008; Gozli, Chasteen, & Pratt, 2013; Ostarek & Vigliocco, 2017). Behavioral work has established that processing up/down words enhances spatial attention to the compatible location which often leads to facilitated detection and identification of visual targets in the primed location (Dudschig, Lachmair, de la Vega, De Filippis, & Kaup, 2012; Gozli et al., 2013; Ostarek & Vigliocco, 2017). Interference in the compatible location is typically observed when stimulus onset asynchrony between prime words and visual targets is short (< 400ms) and when they are semantically unrelated (Estes et al., 2015, 2008; Gozli et al., 2013). Recent eye-tracking studies found that up/down words speed up subsequent spatially compatible saccades (Dudschig et al., 2013; Dunn, 2016) and influence concurrent saccades in the congruent direction (Ostarek, Ishag, Joosen, & Huettig, in press), suggesting that processing these words pre-activates specific motor programs in saccade-related brain areas. However, as these studies relied on paradigms with visual targets in top/down locations, inferences about processes directly activated by the words (in the absence of a visual-spatial task) are limited.

To directly address this question, we assessed whether conceptual processing of up/down words activates direction-specific patterns in the cortical network for saccadic eye movements and spatial attention (Corbetta et al., 1998; Grosbras, Laird, & Paus, 2005; Paus, 1996; Vernet, Quentin, Chanes, Mitsumasu, & Valero-Cabré, 2014). In particular, we predicted that a linear classifier could decode whether a word is ‘up’ or ‘down’ from multi-voxel activity patterns in the frontal eye field in the premotor cortex, the supplementary eye field in the supplementary motor area, the intraparietal sulcus as well as surrounding cortex in the superior and inferior parietal lobules. We further predicted shared activity patterns with up/down saccadic eye movements in these regions. A previous decoding study has reported that spatial judgements about visually presented geometric shapes (slightly above vs. below the centre) and spatial judgements about up/down words (whether the word referent’s typical location is up vs. down) trigger similar patterns in the inferior parietal lobule (Quadflieg et al., 2011). Although this finding is consistent with a shared neural substrate for spatial judgements based on different inputs, it remains unclear whether semantic processing of these words activates direction-specific patterns in the brain networks for spatial attention. Here, we used a task (concreteness judgement) that did not allude to the words’ spatial properties (and this dimension went entirely un-noticed by the participants). Thus, we were able to directly test the involvement of the oculomotor network in semantic processing.

## Methods

### Participants

We recruited 20 native Dutch participants (11 female, mean age: 22) from the local Radboud University database. One participant was excluded because she fell asleep during the experiment and for another participant no data were collected because the scanner stopped working. All analyses we report were based on the remaining 18 participants (10 female, mean age: 21). Participants gave written consent and were paid 12.50 euros. We had ethical approval from the local ethics committee (dossier CMO 2014/288), assigned by the accredited local ethics board CMO Arnhem-Nijmegen.

### MRI Acquisition

Data acquisition was performed on a Siemens Prisma Fit 3T scanner at the Donders Center for Cognitive Neuroimaging using a 32-channel head coil. Functional images were acquired using a multiband sequence with 68 slices, acceleration factor = 4, spatial resolution = 2.0×2.0×2.0 mm, TR = 1.5 s, TE = 39.40 ms. There were two functional runs (ca. 11 minutes each) for the language task and one run for the saccadic eye movement task (ca. 15 minutes). To obtain a high-resolution anatomical image, a t1-weighted MPRAGE sequence was run (192 volumes, spatial resolution = 1mm).

### Stimuli, Procedure, and Design

After receiving task instructions, participants were accompanied into the scanning room where they were asked to put earphones after which their head was stabilised with cushions and the head coil was put in place. Before the experiment started, we made a sound check by playing a word stimulus repeatedly with the scanner noise on, giving the participants an opportunity to adjust the volume until they could hear the words fine. Then, the two language runs were carried out, followed by the saccadic eye movement run, and finally by the anatomical scan.

The main experiment used 24 concrete and 24 abstract filler words. Of the concrete words (see Table 1), 12 referred to objects that are usually observed in the lower visual field (shoe), and 12 referred to objects that are usually up (bird). These two word classes were closely matched for frequency (SUBTLEX: Keuleers, Brysbaert, & New, 2010), age of acquisition (Brysbaert, Stevens, De Deyne, Voorspoels, & Storms, 2014), concreteness (Brysbaert et al., 2014) number of syllables, number of letters, and number of phonemes (all p-values > 0.4). The abstract filler words differed strongly in concreteness (p<0.001), but were similar in frequency, length, number of letters, phonemes, and syllables (p>0.05).

**Table 1:**
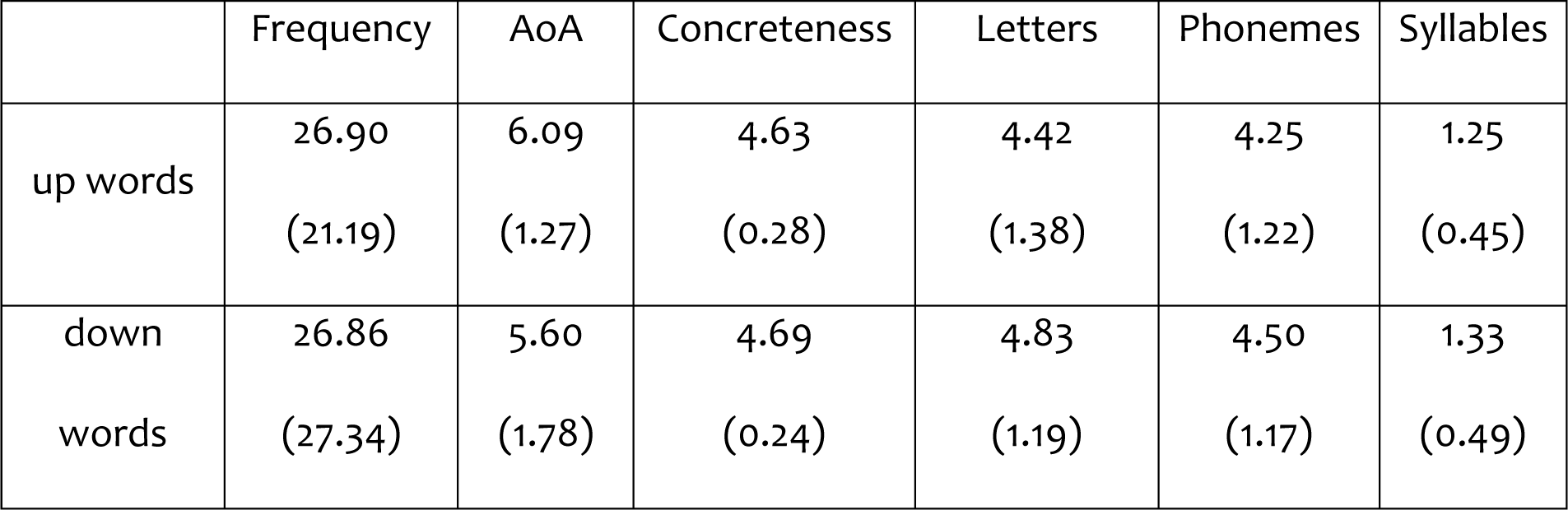
Stimulus characteristics. Means per word category and standard deviations in parentheses. AoA = Age of acquisition.

|  | Frequency | AoA | Concreteness | Letters | Phonemes | Syllables |
| --- | --- | --- | --- | --- | --- | --- |
| up words | 26.90<br>(21.19) | 6.09<br>(1.27) | 4.63<br>(0.28) | 4.42<br>(1.38) | 4.25<br>(1.22) | 1.25<br>(0.45) |
| down words | 26.86<br>(27.34) | 5.60<br>(1.78) | 4.69<br>(0.24) | 4.83<br>(1.19) | 4.50<br>(1.17) | 1.33<br>(0.49) |

A different group of 38 participants were asked to rate the up/down words in terms of whether they refer to objects that are typically perceived up vs. down in space. On a Likert scale ranging from 1=“always down” to 7=“always up”, down words were given a mean rating of 1.84 (SD=1.15) compared to a mean of 6.25 (SD=0.99) for up words (p<0.001).

There was an auditory and a written version of each item (intended to avoid low-level confounds in classification), which were both repeated three times, amounting to six repetitions per word across the whole language experiment and to 288 trials in total (ca. 22 minutes). The order of trials was fully randomized per run. Trials lasted 4.5 seconds (3 TRs). Every spoken word trial began with a spoken word (average: 600ms) accompanied with a fixation dot that remained on the screen until the end of the trial. In written word trials, the written word appeared centrally for 600ms and was then replaced by the fixation dot for the remaining 3900ms. Participants were asked to respond by pressing the correct button (left or right index finger) as quickly as possible without sacrificing accuracy right after word presentation. The task was to decide whether the word was concrete or abstract (concreteness task).

After the language task, participants performed 71 mini-blocks of saccadic eye-movements in which they had to follow a fixation cross with their eyes as closely as possible (the method was based on Knops, Thirion, Hubbard, Michel, & Dehaene, 2009). On every trial, the fixation cross was first presented centrally for 3s and then moved either to the top or bottom of the screen in four steps that were slightly jittered on the vertical and horizontal axis: On the vertical axis the visual angle at each step randomly varied between 1.71-2.14° and the horizontal location was randomly jittered between 0 and 0.64° to the left or right of the centre on the horizontal axis. The speed at which the four steps occurred was varied randomly across mini-blocks (the fixation cross remained at each location for 750ms, 1125ms, or 1500ms). After the fourth step the fixation cross remained at the last position for 6750ms, 5650ms, or 4500ms depending on the speed of the previous steps such that the total time per mini-block was kept constant at 12s.

### Eye-tracking pre-test

As recent studies indicated that processing up/down words can modulate subsequent and concurrent saccades, we conducted a pre-test to rule out that participants move their eyes up/down during central fixation when performing a semantic task on these words. We recruited a separate group of 36 participants for an eye-tracking experiment using an Eye-Link 1000 tracker (SR Research) in which they performed the same task (concreteness judgment) on the same set of words we used in the MRI experiment. Spoken words were presented via headphones and participants were asked to maintain central fixation. A linear mixed effects model was used to analyse possible differences in vertical looking behaviour as a function of the words’ spatial associations. The mode included spatial association (up vs. down) as fixed effect and had per-participant intercepts and slopes. Our results showed that central fixation was maintained remarkably well (up words: mean y-coordinate=5.65) and there was no effect of the spatial association of words on y-coordinates (up words: M=5.65, SD=2.30; down words: M=5.65, SD=2.25; t<1).

### Preprocessing

Preprocessing of the volumes was done using the SPM12 fMRI toolkit for MATLAB. Volumes were slice time corrected to the middle slice and spatially realigned to the first volume of the first run. Volumes were co-registered to the t1-weighted structural scan and the structural scan was segmented by tissue type using SPM12’s segmentation subroutine. Both the functional volumes and the structural scans were normalized to the MNI template.

### General linear model

Functional volumes were entered into a general linear model in which each trial was modelled as a separate regressor (the six motion parameters and per-run parameters were included as regressors of no interest). The regressors corresponding to word events were convolved with the canonical hemodynamic response function and the model parameters were estimated. This procedure yielded 144 beta maps. Beta maps were then aggregated by word and t-value maps were computed (with each t-map being informed by six beta maps of the same word, three of which corresponding to spoken and three to written presentation of the word). This procedure yielded 24 t-maps per run and 48 t-maps in total (i.e. one per item).

### Searchlight analysis

From the 48 word t-maps, the 24 maps associated with concrete words were selected, discarding the 24 maps associated with abstract words. The 24 concrete word maps were further split into a group of 12 maps associated with up words and a group of 12 maps associated with down words.

To identify brain regions whose patterns distinguish up words from down words, a whole-brain searchlight procedure was used, as proposed by Kriegeskorte, Goebel, and Bandettini (2006) and implemented in the PyMVPA toolkit (Hanke et al., 2009). For each voxel in the map, a sphere with a radius of 3 voxels around that voxel was selected and a linear kernel support vector machine classifier was trained on that sphere of voxels in twenty of the t-maps and then tested on the four remaining t-maps. This procedure was cross-validated by repeating it on 10 different balanced folds of the dataset per sphere to ensure a stable estimate of prediction accuracy.

To identify clusters with significant prediction accuracy, a combination of training set permutation and bootstrapping followed by cluster-size control and correction for multiple testing was used, as proposed by Stelzer, Chen, and Turner (2013), as implemented in the PyMVPA toolkit (Hanke et al., 2009): To allow statistical inferences based on the searchlight prediction accuracy, an empirical null distribution was estimated based on a permutation-bootstrapping scheme. In the permutation step, a chance accuracy map was generated by repeating the searchlight and cross-validation procedure with randomly shuffled t-map labels. This procedure was repeated 100 times per participant. A group-level null distribution was then estimated using a bootstrap procedure by selecting one chance accuracy map per participant and computing an averaged group chance map 100,000 times.

Statistically significant clusters were identified by first clustering contiguous voxels significant at the .001-level, computing the cluster-wise p-value and then correcting these p-values using the Benjamini-Hochberg method for false discovery rate correction (Benjamini & Hochberg, 1995).

To assess whether brain activity patterns were shared between looking up vs. down and processing an up vs. down word, we performed a cross-classification analysis. Cross-decoding from saccades (upwards/downwards) to words (up/down) was done by performing a searchlight analysis (radius = 3 voxels) in which a linear support vector machine classifier was trained on the patterns related to the saccadic eye-movement data and then tested on the patterns corresponding to the 24 up/down words. To ensure stable estimates, for every searchlight we used a cross-validation scheme where the classifier was trained on all but six randomly chosen saccade trials and tested on all up/down words. This procedure was repeated 10 times and the average accuracy across the 10 folds was taken as the central voxel’s cross-decoding accuracy value. Cluster-based permutation testing was performed by again repeating this analysis 100 times per participant with randomly shuffled labels in the training set to then estimate a group-level null distribution as described above for up/down word decoding.

## Results

### Behavioral results

Participants performed the concreteness task accurately and, importantly, achieved highly similar accuracies for up words (93.85%) and down words (94.01%) that were statistically indistinguishable (t<1). Reaction times to up words (M=1105 ms, SD=435 ms) and down words (M=1123 ms, SD=462 ms) were also very similar (t<1).

### MVPA Results

As a first step, we used searchlight analysis to identify brain regions whose multi-voxel patterns can discriminate up from down words (see Figure 1). Cluster-based permutation tests revealed successful decoding in a set of regions linked to saccadic eye-movements and spatial attention, including the posterior portion of the superior frontal sulcus, the middle frontal gyrus, the inferior and superior parietal lobule, posterior temporal lobe, the cerebellum, and portions of the occipital cortex including primary visual cortex.

**Figure 1:**
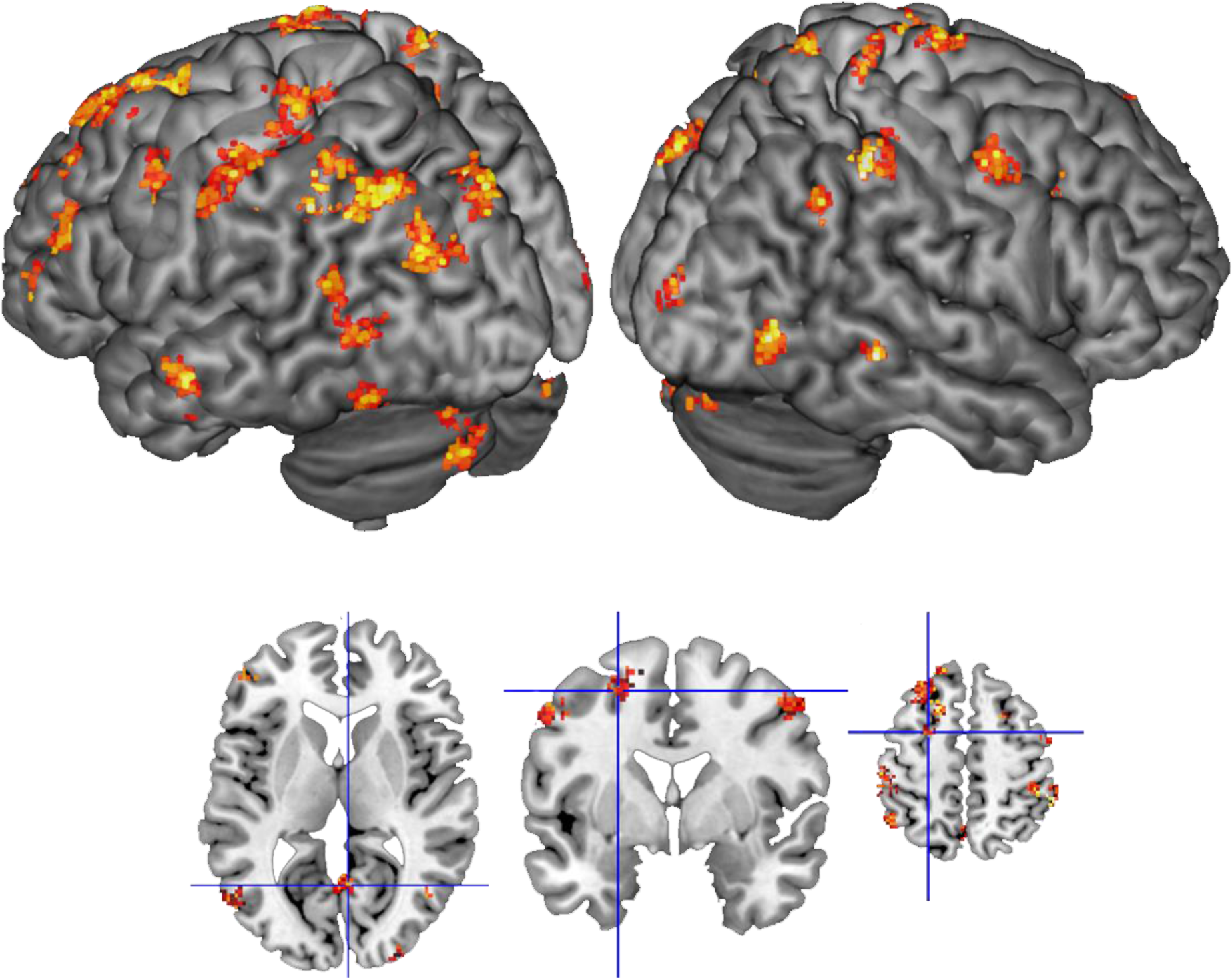
Up/down word decoding results. Top: Significant clusters on the cortical surface. Bottom (from left to right): Axial slice with crosshairs on the right primary visual cortex; coronal and axial slice with crosshairs on the posterior superior frontal sulcus.

To provide direct evidence that up/down words engage processes similar to actual up/down shifts of attention, we sought to train a classifier to distinguish up vs. down saccades and tested it on the up/down words without further training. Decoding saccade direction is expected to be possible based on activity patterns in a network of occipital, parietal, and frontal areas. However, we obtained above-chance decoding in virtually the entire brain and there was a prominent peak with decoding accuracies close to 1 in the white matter behind the eyes. This suggests that the eye movements induced small motion artefacts. We carried out cross-decoding nevertheless based on the following grounds: 1) Peaks in saccade decoding (looking up vs. down) were observed in the occipital cortex, posterior parietal cortex, and the frontal eye field region at the intersection of the superior frontal sulcus and the precentral sulcus, as expected (Figure 2 shows regions with decoding accuracies higher than 60%). 2) It seems implausible that motion artefacts induce fine-grained activation patterns that resemble the ones evoked by up/down words. Instead, generalization of patterns from the saccade to the word domain should only succeed based on brain activity related to saccade planning and execution.

**Figure 2:**
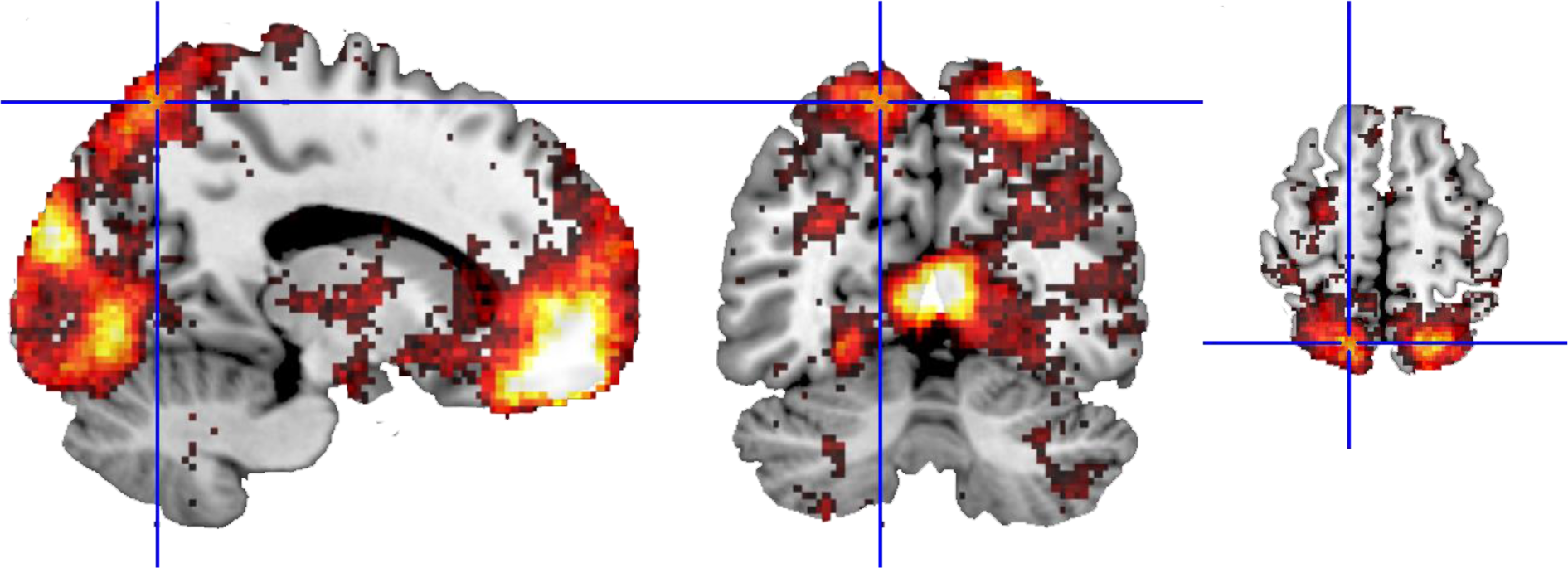
Saccade direction decoding. Sagittal (left), coronal (middle), and axial (right) slices plotting decoding accuracies > 60%.

In line with this rationale, no significant cross-decoding was observed in the white matter behind the eyes that appeared to be most susceptible to motion artefacts. Instead, the cross-decoding approach revealed very robust clusters in the intraparietal sulcus, superior parietal lobule, inferior parietal lobule, the intersection of the superior frontal sulcus and the precentral sulcus (FEF), supplementary motor area, prefrontal cortex, precuneus, posterior middle and superior temporal lobe, posterior inferior temporal cortex, cingulate gyrus, and in the occipital lobe including primary visual cortex (see Figure 3).

**Figure 3:**
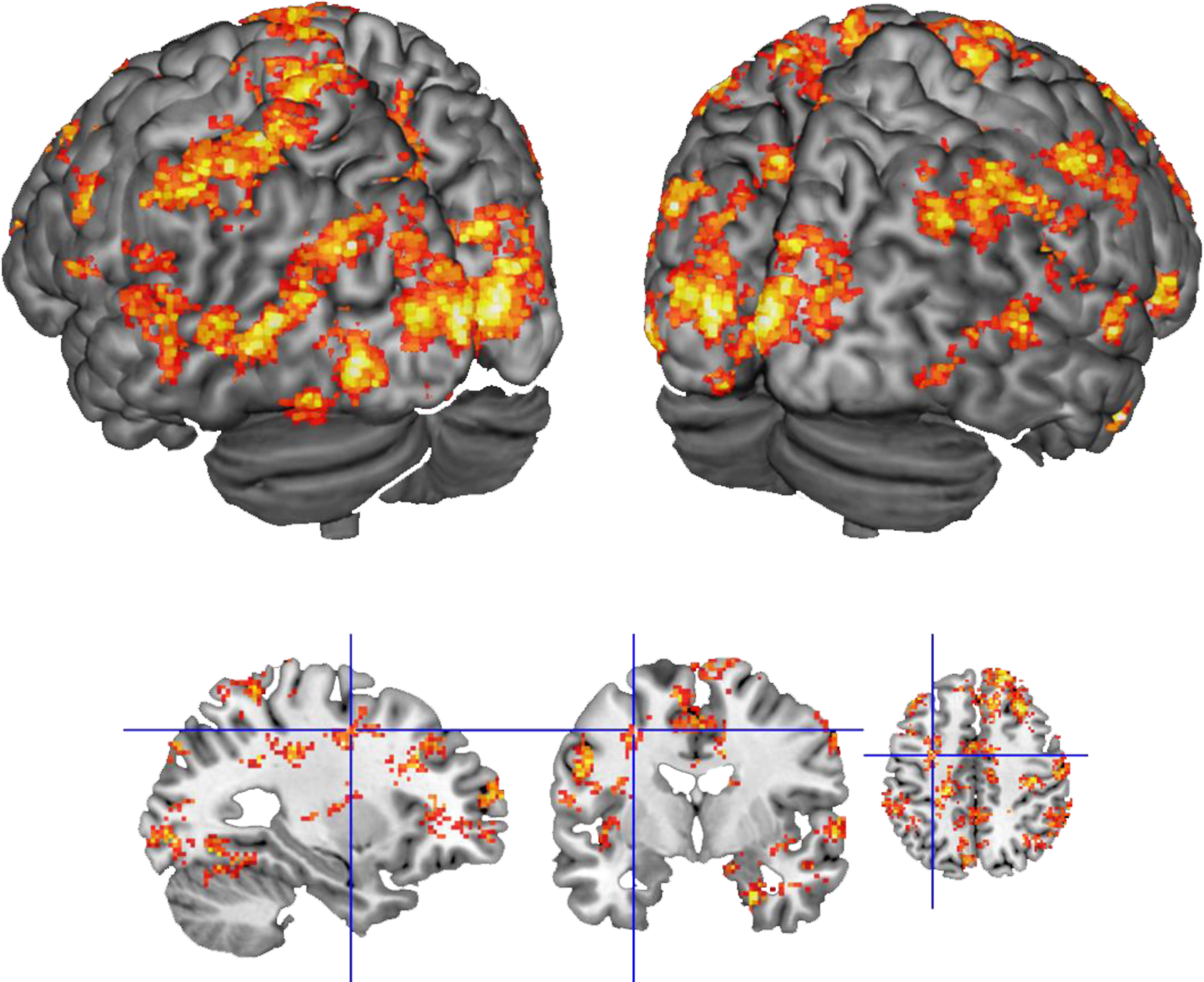
Cross-decoding results. Top: Significant cross-decoding from saccades to words in visual, temporal, and parietal regions. Bottom (left to right): Sagittal, coronal, and axial slices with crosshairs on the FEF.

Overlap between word decoding and cross-decoding (Figure 4) was observed in the inferior parietal lobule (angular gyrus and left supramarginal gyrus), precuneus, middle and superior occipital gyrus, posterior middle temporal gyrus, supplementary motor area, superior frontal gyrus, and superior frontal sulcus towards the precentral sulcus.

**Figure 4:**
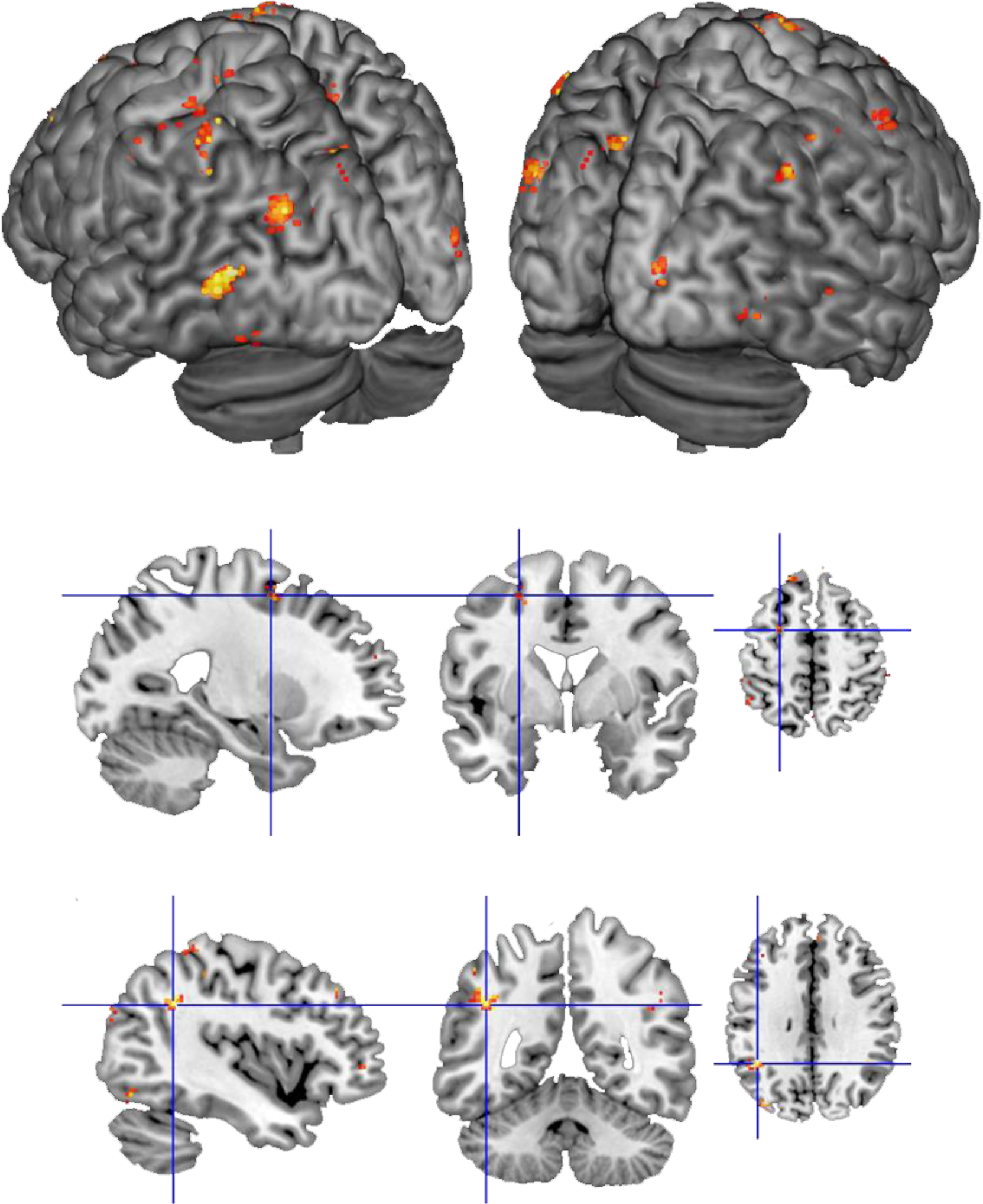
Overlap between cross-decoding and word decoding. The cortical surface mainly shows overlap in the middle occipital gyri, posterior inferior and middle temporal lobes (top panel). Overlap was also observed in the posterior superior frontal sulcus (middle panel) and the inferior parietal lobule (bottom panel).

## Discussion

Previous behavioural and eye-tracking studies suggested that words with implicit spatial (up/down) associations can subtly influence performance on simple visual identification tasks involving up/down targets (Gozli et al., 2013). Up/down words have further been reported to increase the speed of subsequent saccadic eye movement initiation (Dudschig et al., 2013; Dunn, 2016), and to modulate the trajectories of concurrently performed saccades (Ostarek et al., in press). Here, we set out to test the hypothesis that semantic processing of up/down words in the absence of eye movements recruits the oculomotor system. In line with this hypothesis, we found that multi-voxel patterns in several key areas implicated in programming and executing eye movements allowed reliable decoding of up vs. down words and cross-decoding from saccades to words.

Two related lines of research make this finding particularly interesting. The first capitalizes on the insight that covert shifts of spatial attention have a largely shared neural substrate with covert spatial attention (Corbetta et al., 1998; Fan, McCandliss, Fossella, Flombaum, & Posner, 2005; Grosbras et al., 2005). This can be accounted for by the premotor theory of attention (Rizzolatti, Riggio, Dascola, & Umiltá, 1987) holding that the same neurons are active for saccades and covert shifts of attention, or by models assuming overlapping but distinct interacting cell populations (Petersen & Posner, 2012; Schafer & Moore, 2007; Thompson, Biscoe, & Sato, 2005). Distinguishing between the two is not possible in fMRI because of limitations in spatial resolution, but we can conclude that on a systems level, similarly to covert shifts of attention, semantic information about the typical location of an item is retrieved from the cortical network for goal-directed eye movements.

The second line of research is the work on the mental number line and the neural correlates of arithmetic processing that has established a strong link with the cortical systems for orientation in horizontal space. For instance, one study reported successful cross-decoding from left vs. right saccades to subtraction vs. addition (Knops et al., 2009). Furthermore, when trying to bisect numerical intervals patients with left neglect show an error pattern with a shift towards larger numbers (Zorzi, Priftis, & Umiltà, 2002), similar to the bisection of physical lines where a shift to the right relative to the midpoint was observed (Schenkenberg, Bradford, & Ajax, 1980). This had led to the view that cultural acquisitions, such as numerical cognition, recycle evolutionary older systems that perform useful computations for the new task at hand (Dehaene, 2005; Dehaene & Cohen, 2007).

In the context of these considerations, it thus seems reasonable to propose that a similar phenomenon occurs in the domain of language comprehension. In particular, our results indicate that the dorsal-frontal system for overt and covert shifts of attention is recycled to provide semantic information about the typical vertical spatial location of word referents. Previous studies have shown that explicit language about spatial relationships (Kemmerer & Tranel, 2003; Tranel & Kemmerer, 2004) and judgments about word referents’ typical location (Quadflieg et al., 2011) recruit parietal areas involved in spatial cognition. The novel contribution of our data is that they implicate the orienting network with conceptual processing of implicitly spatial words in a semantic task orthogonal to the spatial domain.

Besides the spatial processing/oculomotor areas, information about typical location was present in temporal and prefrontal areas that are part of the ‘core’ semantic network (Binder et al., 2009). This suggests that access to semantic information about space relies on interactions between high-level areas typically associated with lexical-semantic processing on the one hand, and spatial processing/oculomotor areas on the other. Future studies benefiting from higher temporal resolution or using functional connectivity analysis could investigate the flow of information between these systems.

## Acknowledgments

We would like to thank Paul Gaalman and José Marques for their help with the preparation of the MRI sequences and data collection.

